# An asexual flower of *Silene latifolia* and *Microbotryum lychnidis-dioicae* promoting its sexual-organ development

**DOI:** 10.1101/634725

**Authors:** Hiroki Kawamoto, Kaori Yamanaka, Ayako koizumi, Kotaro Ishii, Yusuke Kazama, Tomoko Abe, Shigeyuki Kawano

**Author notes:** Functional Biotechnology PJ, Future Center Initiative, The University of Tokyo, Wakashiba 178-4-4, Kashiwa, Chiba 277-0871, Japan. **Corresponding author:** Shigeyuki Kawano.

## Abstract

*Silene latifolia* is a dioecious flowering plant with sex chromosomes in the family Caryophyllaceae. Development of a gynoecium and stamens are suppressed in the male and female flowers of *S. latifolia*, respectively. *Microbtryum lychnidis-dioicae* promotes stamen development when it infects the female flower. If suppression of the stamen and gynoecium development is regulated by the same mechanism, suppression of gynoecium and stamen development is released simultaneously with the infection by *M. lychnidis-dioicae*. To assess this hypothesis, an asexual mutant, without gynoecium or stamen, was infected with *M. lychnidis-dioicae*. A filament of the stamen in the infected asexual mutant was elongated at stages 11 and 12 of the flower bud development as well as the male, but the gynoecium did not form. Instead of the gynoecium, a filamentous structure was suppressed as in the male flower. Developmental suppression of the stamen was released by *M. lychnidis-dioicae*, but that of gynoecium development was not released. It is thought, therefore, that the suppression of gynoecium development was not released by the infection of *M. lychnidis-dioicae. M. lychnidis-dioicae* would have a function similar to SPF since the elongation of the stamen that is not observed in the healthy asexual mutant was observed after stage 8 of flower bud development. Such an infection experiment also that the Y chromosome of the asexual mutant has genes related to the differentiation of archesporial cells, but none related to maturation of the tapetal cells.

## Introduction

The basidiomycetous genus *Microbotryum* contains a species-rich member of smut fungi that infects a wide range of the host plant belonging to Caryophyllaceae, Dipsacaceae, Lamiaceae, and Lentibulariaceae in the dicotyledonous plants [1]. The smut fungus *M. lychnidis-dioicae* was isolated from *S. latifolia* [2]. *M. lychnidis-dioicae* has long been used as a model for the study of ecology, genetics, and the propagation of sexual infection [3], [4], [5]. Smut disease is transmitted by smut spores attached to the insect [6]. Spore germination, meiosis, and mating of paired basidiospores of smut fungi occurs on the host cell surface [7]. The basidiospores of smut fungi form a secondary mycelium after mating, which penetrates into the host plant. The secondary mycelium of smut fungi is observed in intercellular regions, vascular bundles, and apical meristem after invasion of the host plant. Subsequently, the secondary mycelia of smut fungi form smut spores in the pollen sacs at the blossom of host plants [8].

*S. latifolia* is a dioecious flowering plant with sex chromosomes in the family Caryophyllaceae; individual plants have male and female flowers. The female flowers blossom with 22 autosomes and two X chromosomes, and the male flowers blossom with 22 autosomes, one X chromosome, and one Y chromosome. It is known that *S. latifolia* is a model for the study of the evolution of plant sex chromosomes and ecology [9]. We require sex-chromosome-linked markers to better understand plant sex chromosomes. Therefore, several Y chromosome-linked markers were made using Amplified Fragment Length Polymorphism (AFLP) [10], Random Amplified Polymorphic DNA (RAPD) [11], and a technique that combines laser microdissection and polymerase chain reaction (PCR) [12]. The male flowers of *S. latifolia* with Y chromosomes were irradiated with γ rays or heavy ion beams to produce hermaphrodites, an asexual mutant, and a pollen-defect mutant with deletions to a part of the Y chromosome [13], [14], [15]. Using the deletion status of these mutants, a map of the *S. latifolia* Y chromosome was created [10], [16], [17], [18], [19], [20], [21]. Phenotypes differ between smut infected-male and female hosts. *M. lychnidis-dioicae* forms a smut spore instead of pollen in the pollen sac in the infected male. In contrast, stamen formation is promoted in the infected female. Smut spores are then formed instead of the pollen in anthers [22], [23].

It is thought that three male factors: gynoecium suppression factor (GSF), stamen promoting factor (SPF), and male fertility factor (MFF), exist on the Y chromosome of *S. latifolia*. The asexual mutant was first isolated by Donnison et al. [14]. The asexual mutant has X and Y chromosomes, but the SPF region is deleted on the Y chromosome. The flowers of asexual mutants have a filamentous structure instead of the gynoecium in the center of the flower (as observed in males), as well as developmentally suppressed stamens at the early developmental stages [14], [24]. Sporogenous cells in developmentally suppressed anthers of the asexual mutant form in the flower buds at very early developmental stages, but parietal cell layers are absent [25].

What happens when *M. lychnidis-dioicae* infects asexual mutants with parietal deletions in the Y chromosome? In asexual mutants, development of the stamen and gynoecium is suppressed. If suppression of stamen and gynoecium development results from the same mechanism, then suppression of gynoecium development should be released when suppression of stamen development is released by *M. lychnidis-dioicae* infection. In addition, what differences exist in stamens between the infected asexual mutant, the infected male, and the infected female? We infer that we are able to search for genes related to the anther development, which should exist in the deleted region of the Y chromosome, by comparing pollen sacs of stamens in the infected male, infected female, and infected asexual mutant.

In this study, progeny of the asexual mutant could be successively produced by crossing the female-like flowers in the asexual mutant with the male flower in the wild-type male [24]. The infected asexual mutant was found as 5 individuals due to inoculation with M. lychnidis-dioicae and PCR screening. We especially focus on gynoecium development in the infected asexual mutant and successively compared the morphological changes in floral organs caused by all *M. lychnidis-dioicae* infection among males, females, and asexual mutants using scanning electron microscopy and tissue section analysis.

## Materials and Methods

### Plant materials and plant growth conditions

*S. latifolia* seeds were obtained from the inbred line (K-line) and stored in our laboratory. The K-line was propagated for 17 generations of inbreeding to obtain a genetically homogeneous population. We also used asexual mutants obtained from crossing the inbred K-line with an asexual mutant (ESS1), which was originally one of heavy-ion beam irradiation-induced Y-deletion mutants identified by Fujita et al. [20]. Plants were grown from vernalized seeds in pots in a regulated chamber at 23°C with a 16-h light/8-h dark cycle.

### M. lychnidis-dioicae *inoculation*

Sporidia of A1 and A2 were cultured on potato dextrose agar (BD Difco) at 23°C for 5 days and suspended at 2 × 10^6^ cells/mL in distilled water. Sporidial mixtures of A1 and A2 at equal concentrations were used throughout the inoculations. Inoculation treatments were carried out on 10-d-seedlings of *S. latifolia* on 0.8% agar plates. The base of each 10-d-old seedling was injected with 2 μL of the mixture. Inoculation was repeated after 3 days. Three weeks after inoculation, we transferred seedlings to soil in pots and grew them in a regulated chamber at 23°C with a 16-h light/8-h dark cycle.

### Scanning electron microscopy

Flowers were fixed overnight in 2.5% glutaraldehyde in 0.1 M phosphate buffer (pH 7.2) at 4°C. After being washed three times with 0.1 M phosphate buffer (pH 7.2), fixed flowers were dehydrated in an ethanol series (30, 50, 70, 80, 90, 95, and 100% each step for 30 min at room temperature) and kept in 100% ethanol overnight at 4°C. The ethanol was replaced with isopentyl acetate and the flowers were dried with a critical-point dryer (HCP-2, Hitachi, Tokyo) and sputter-coated with platinum palladium using an ion sputter (E-1010, Hitachi). The flowers were examined in an S-3000N scanning electron microscope (SEM) (Hitachi) operated at 5 kV in the high vacuum mode. The gynoecium on the scanning microphotographs was colored with Photoshop 7.0 (Adobe Systems, Inc., San Jose, CA).

### Light microscopy

Flowers were double fixed overnight in 4% glutaraldehyde in 0.1 M phosphate buffer (pH 7.2) at 4°C and post-fixed for four hours in 2% osmium tetraoxide in distilled water. After being washed with 0.1 M phosphate buffer (pH 7.2), the fixed flowers were dehydrated in an ethanol series (30, 50, 70, 80, 90, 95, and 100% each step for 15 min at room temperature) and kept in 100% ethanol overnight at 4°C. The ethanol was replaced with xylene and embedded in paraffin. Embedded flowers in paraffin were cut into 10-μm sections using a microtome (RV-240, Yamato, Japan). Cutting sections were de-paraffinized in xylene and rehydrated in an ethanol series (100, 95, 90, 80, 70, 50, and 30%; each step for 10 min at room temperature). Rehydrated sections were stained Schiff’s reagent. The sections were observed with a microscope (BX60, Olympus, Tokyo, Japan).

### Screening for asexual mutants with partial Y chromosome deletion

To obtain asexual mutants from crossing a sexual mutant (ESS1) and inbred line (K-line), we performed polymerase chain reaction (PCR) screening using four STS makers (MS4, MK17, ScQ14, and SlAP3) and flower phenotype screening. STS makers (MS4, MK17, ScQ14, and SlAP3) are described in the kazama et al., [21]. Genomic DNA was extracted from fresh leaves using a DNeasy Plant Mini Kit (Qiagen) according to the manufacturer’s instructions. PCR amplification was performed using Blend Taq polymerase (Toyobo, Tokyo, Japan) with a Thermal Cycler Dice TP600 (TaKaRa Bio, Otsu, Japan). The conditions were 5 min at 94°C, followed by 35 cycles of 1 min at 94°C, 1 min at 60°C, and 1 min at 72°C, with final extension of 5 min at 72°C. Each 20-μL reaction mixture contained 50 ng of template DNA, 2 μL dNTPs, 10 × Blend Taq Buffer, 0.2 μL of Blend Taq, and 1 μL of each primer at 5 mM. The reaction products were electrophoresed on 1.5% agarose gels. After staining with ethidium bromide, fragments were visualized on a UV illuminator (Atto, Tokyo, Japan). Wild-type male and female genomic DNA was used as a control.

## Results

### Stages of flower bud development in a S. latifolia asexual mutant

In this study, we newly defined stages of flower bud development of the asexual mutant to correspond with stages of flower bud development in male and female flowers. Stages of flower bud development of the asexual mutant were divided into 11 stages (Fig. 1a–k). Flower bud development in the asexual mutant at stages 1 to 6 was similar to that of males. The flowers of the asexual mutant with developing stamen primordia and gynoecium primordia at stages 1 to 5 of flower bud development showed hermaphroditic morphology (Fig. 1a–f). However, although esxpansion of the stamen filament occurred at stage 7 in male flowers [23], **(S1 Fig.)**, expansion of the stamen filament was not seen in the asexual mutant at stage 7, (Fig. 1g). Stamen development was suppressed at stage 8 of flower bud development, and the gynoecium became a filamentous structure (Fig. 1h). Only the petals and filamentous gynoecium developed at stages 9 to 11 (Fig. 1i–k). Suppressed anthers became trapezoidal or rectangular, similar to structures observed in the wild-type female (Fig. 1).

**Fig. 1.**
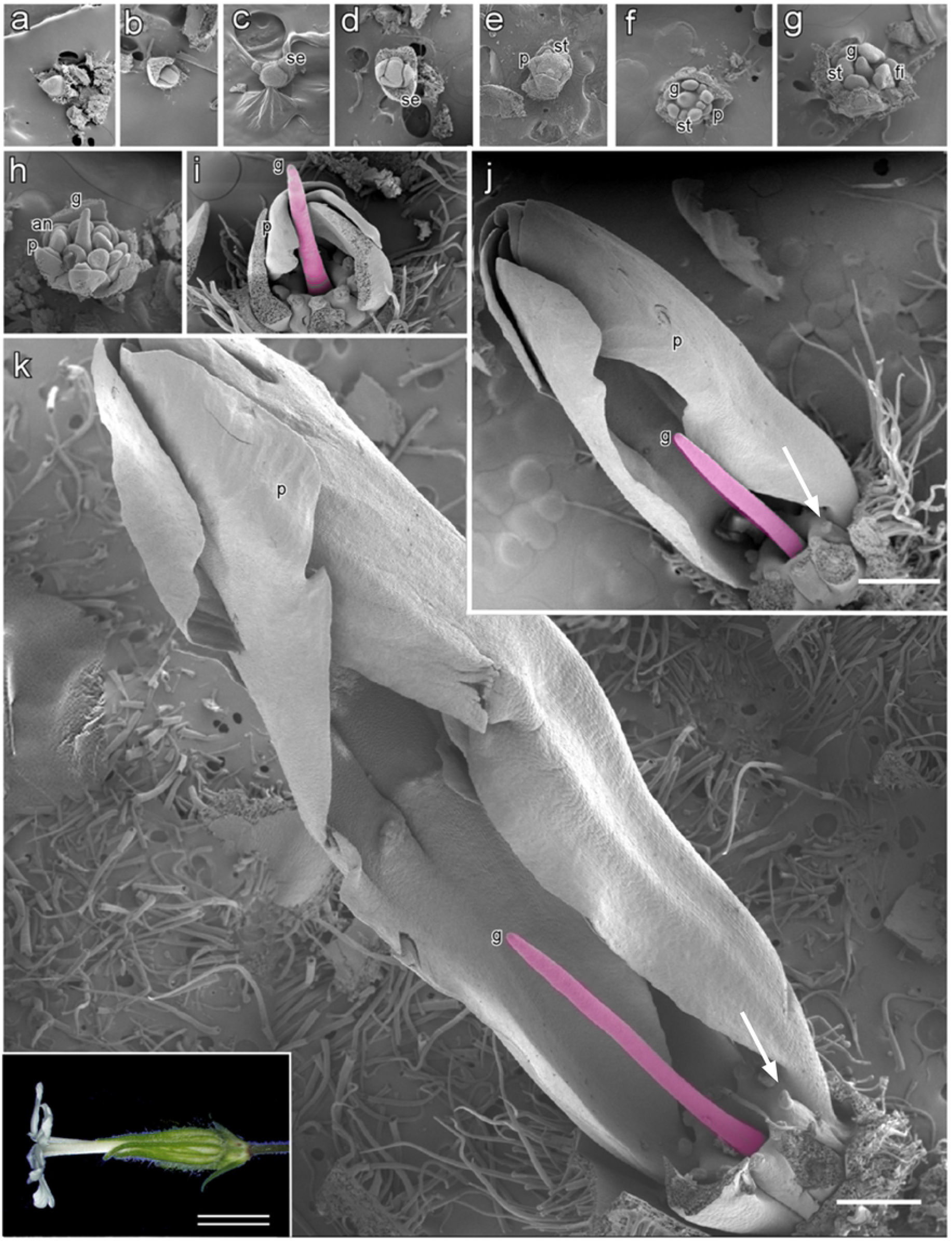
Scanning electron microphotographs of flower bud development in an asexual mutant of *S. latifolia*. Asexual flower buds are shown partially dissected to reveal internal structures at successive stages of development. Developmental stages of the asexual flowers are divided into 11 stages in this study **(a–k)**. Stages 1 to 7 are shown in transverse view; remaining stages are shown in longitudinal view. Early stages are observed to be bisexual **(a–f)**, but later stages are not observed in gynoecium or anther development **(g–k)**. **a** Asexual flowers at stage 1; **b** asexual flowers at stage 2; **c** asexual flowers at stage 3; **d** asexual flowers at stage 4; **e** asexual flowers at stage 5; **f** asexual flowers at stage 6; **g** asexual flowers at stage 7; **h** asexual flowers at stage 8; **i** asexual flowers at stage 9; **j** asexual flowers at stage 10; **k** asexual flowers at stage 11. The inset bright field microphotograph in **k** shows an open flower. Anthers **(an)**, sepal **(se)**, stamen **(st)**, gynoecium **(g)**, petal **(p)**, and filaments **(f).** Arrows in j and k indicate suppressed anthers. Bar = 500 μm, double bar = 1 cm

### Morphological changes in the asexual mutant caused by M. lychnidis-dioicae

Both the asexual flower and the female-like flower blossom are produced in the asexual mutant ESS1, which has one X chromosome and one Y chromosome with a deletion of the SPF region. The female flowers had five styles, but the female-like flowers had only two styles. It was possible to produce progeny of the asexual mutant by crossing the female-like flowers in the asexual mutant with the male flower in the wild-type male because gynoecia of the female-like flowers in the asexual mutant were fertile. As a result, we obtained 234 seeds, and those seeds were sowed. Of the germinated seeds, 188 S. latifolia sprouts were inoculated with *M. lychnidis-dioicae.* We genotyped mock or infected plants three months after infection, based on genotyping by PCR using four markers: MK17, ScQ14, SlAP3, and MS4 (Fig. 2).

**Fig. 2.**
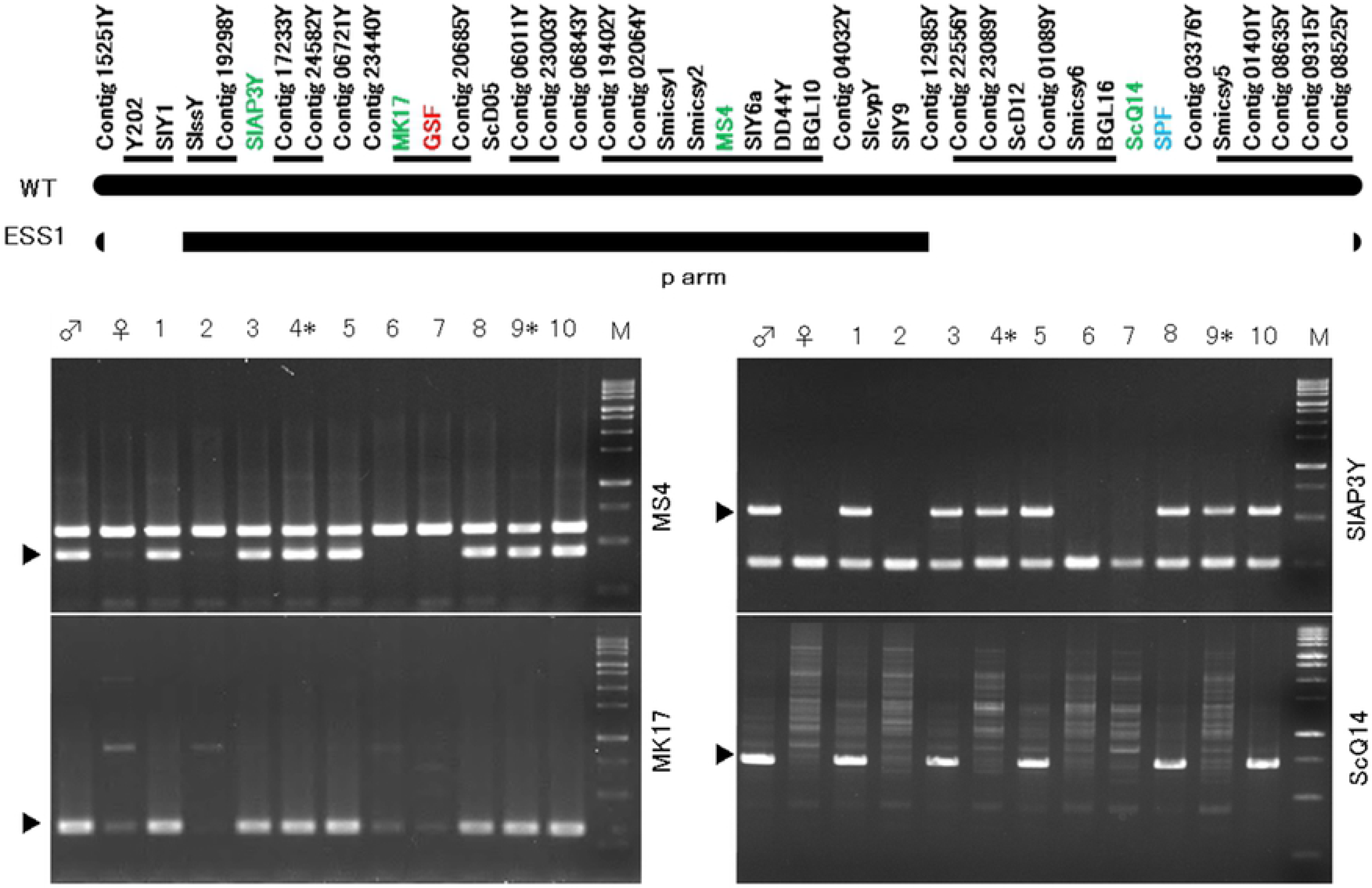
Selection of infected asexual mutant. After that 188 of germinated seeds were inoculated with *M. lychnidis-dioicae* to a sprout of *S. latifolia*. We checked mock or infected after the three months, and those were performed genotyping by PCR using four markers MK17, ScQ14, SlAP3, MS4. asterisk indicate infected asexual mutant. Black arrow head indicate Y-specific band. The order of the markers is as described in Kazama et al. [21]. Closely linked marker sets, in which the orders of markers are not fixed, are underlined.

We obtained 66 female individuals with two X chromosomes, 91 male individuals with the X chromosome and the intact Y chromosome, and 15 asexual mutant individuals with the X chromosome and the Y chromosome with deletion of the SPF region (Table 1). In these plants, the infected female was found as 48 individuals, the infected male was found as 54 individuals, and the infected asexual mutant was found as 5 individuals (Table 1). Flower bud development in the infected asexual mutant was divided into 12 stages (Fig. 3 a–l). The morphology of the infected asexual mutant appeared to be similar to that of the male and healthy asexual mutant at stages 1 to 6 (Fig. 3 a–f). However, extension of the stamen filament in the infected asexual mutant was confirmed, as well as in the male, at stages 7 and 8 of flower bud development, and developmental suppression of the stamens did not occur at stage 8 in the asexual mutant (Fig. 3 g, h).

**Table 1.** Sexual genotype and *M. lychnidis-dioicae* infection ratio in progeny by the cross between asexual mutant (♀) × wild-type male (♂) of *S. latifolia*.

| Phenotypes | Genotypes | Result of inoculations |  | Total |
| --- | --- | --- | --- | --- |
|  |  | Mock | Infected |  |
| Female | XX | 35 | 48 | 83 |
| Male | XY | 36 | 54 | 90 |
| Asexual | XY<sup>d</sup> | 10 | 5 | 15 |
| Total |  | 71 | 107 | 188 |

Y<sup>d</sup> is indicated Y chromosome with deletion.

**Fig. 3.**
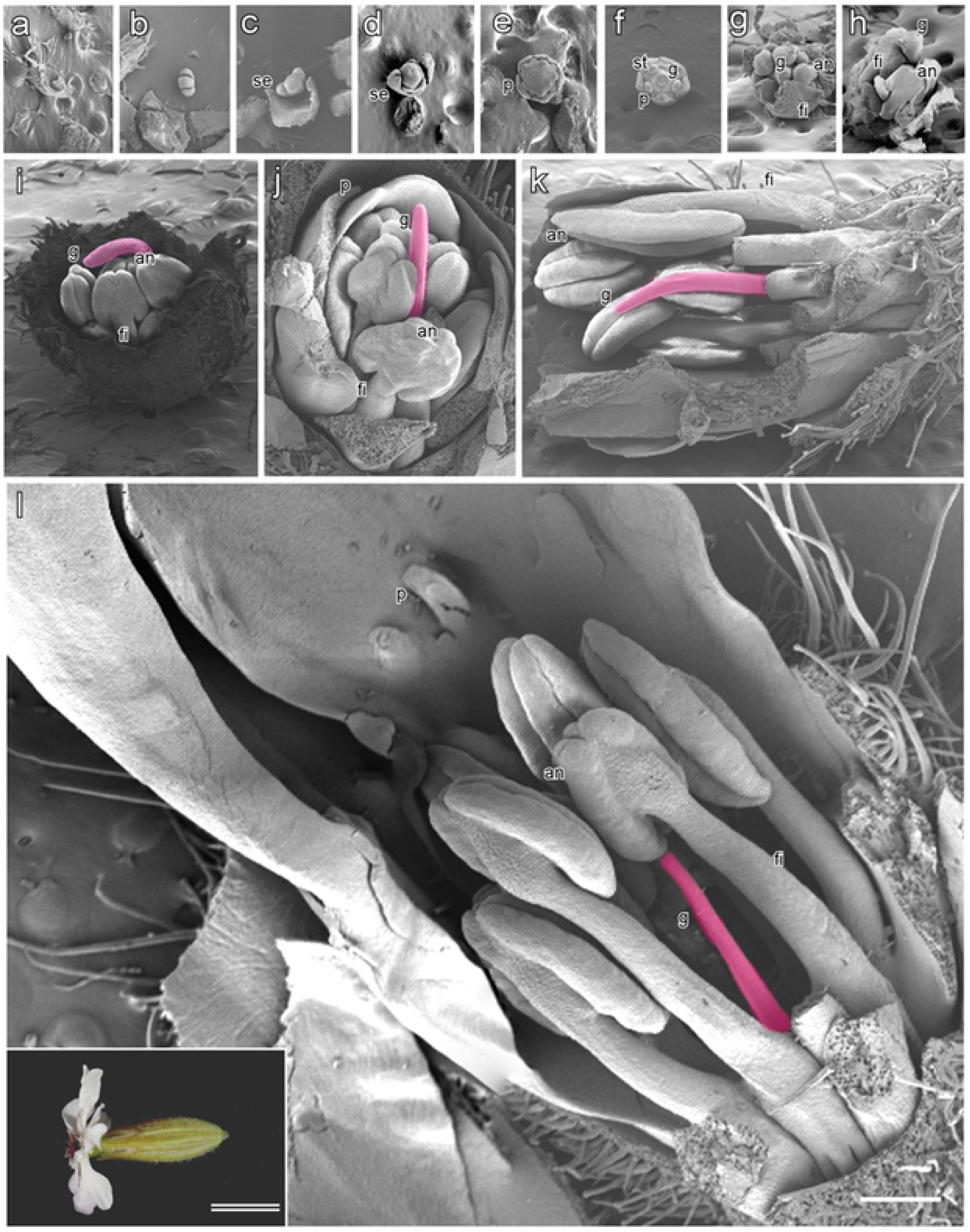
Scanning electron photomicrographs of flower bud development in infected asexual *S. latifolia*. Infected asexual flower buds are shown partially dissected to reveal internal structures at successive stages of development. Developmental stages of the infected asexual flowers are divided into 11 stages in this study **(a–n)**. Stages 1 to 7 are shown in transverse view; remaining stages are shown in longitudinal view. Early stages are observed to be bisexual **(a–f)**, and later stages are anther development, which does not occur in asexual mutants **(g–l)**. **a** Infected asexual flowers at stage 1; **b** infected asexual flowers at stage 2; **c** infected asexual flowers at stage 3; d infected asexual flowers at stage 4; e infected asexual flowers at stage 5; **f** infected asexual flowers at stage 6; **g** infected asexual flowers at stage 7; **h** infected asexual flowers at stage 8; **i** infected asexual anthers at stage 9; **j** infected asexual anthers at stage 10; **k** infected asexual flowers at stage 11; **l** infected asexual flowers at stage 12. The inset bright field microphotograph in **l** shows an open flower. Anthers **(an)**, sepal **(se)**, stamen **(st)**, gynoecium **(g)**, petal **(p)**, and filaments **(f)**; Bar = 500 μm, double bar = 1 cm.

The filamentous gynoecium was extended only in the upward direction and became thinner at stages 8 to 10. The anther in stamens of the infected asexual mutant had four anther locules, as well as the wild-type male (Fig. 3 h–j). The stamen filament of the infected asexual mutant was extended at stages 11 and 12 of flower bud development, as well as the male (Fig. 3 k–l), but anthers were filled by smut spores instead of pollen. In addition, the gynoecium of the infected asexual mutant did not develop. Instead, the filamentous structure formed in the center of the flowers as in the male flowers (Fig. 3 g–l). It was found that developmental suppression of the stamen was released by *M. lychnidis-dioicae*, but that of the gynoecium was not released.

### Morphological changes of anther locule caused by M. lychnidis-dioicae

In this study, we observed anther locules of the healthy male, the infected male, as well as those of the infected female and infected asexual mutants (Fig. 3, Fig. 4 a–y; **S1 Fig.**). Each of the infected plants was observed and compared at stages II to VI of anther development. Archesporial cells and epidermal cells formed at stage II of anther development in the healthy male (Fig. 4 a). The anther locule at stage V of anther development was composed of a layer of epidermis, a layer of endothecium, a middle layer, a tapetum layer and pollen mother cells as a result of the differentiation of archesporial cells (Fig. 4 a–d, f–i). The middle layer and the tapetum layer caused programmed cell death at stage VI of anther development (Fig. 4 e, j).

In the infected males, the primary parietal cell layer was divided into two layers at stage III of anther development (Fig. 4 l). Furthermore, the secondary parietal cell layer was divided into two layers at stage IV of the anther development (Fig. 4 m). In the infected males, the endothecium was present in the two layers, unlike in the healthy males. Furthermore, the tapetum and the middle layer in infected males showed different morphology from those in the healthy males at stage V of anther development (Fig. 4 n). The tapetum was rapidly disintegrated at stage VI of anther development in the infected male, unlike in the healthy male, where the hyphae of *M. lychnidis-dioicae* grew (Fig. 4 o).

In the infected female, only archesporial-like-cells and epidermal cells existed, but primary parietal cells and primary sporogenous cells did not exist in stage III of anther development (Fig. 4 q). Differentiation of the archesporial-like-cells did not occur at later stages (Fig. 4 p–s). Growth of *M. lychnidis-dioicae* was also observed until stage VI of anther development in the infected female (Fig. 4 t). In the infected asexual mutant, development of the anther locules in developed stamens caused by *M. lychnidis-dioicae* was similar to that in the infected male at stages II to IV of anther development (Fig. 4 u–w), whereas the morphology of the tapetum at stage V of anther development was different from that in the infected male (Fig. 4 x). Growth of *M. lychnidis-dioicae* was also observed at stage VI of anther development (Fig. 4 y). Thus, it was found that *M. lychnidis-dioicae* in the infected asexual mutant and infected female grew at the center of the anther locule, where the tapetum was absent, although *M. lychnidis-dioicae* in the infected male grew along the tapetum (Fig. 4 n, s, x; Fig. 5).

**Fig. 4.**
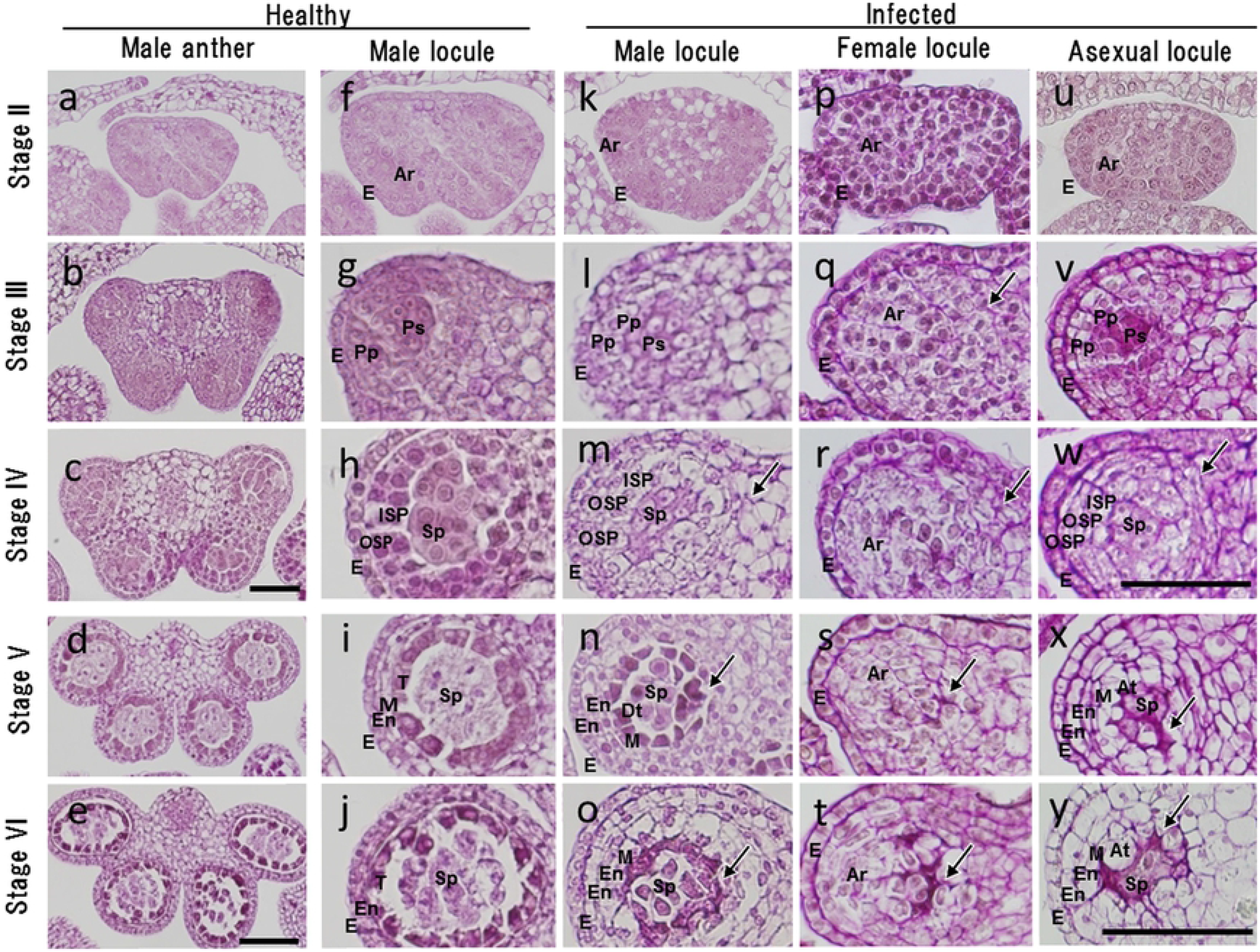
Light photomicrographs showing anther development at stages II to VI in the healthy male flower, the infected male flower, the infected female flower and the infected asexual flower. Developmental stages of healthy male flowers are divided into 14 stages in this study. Four stages of anther development in healthy male flowers and the corresponding stages of development in infected male flowers, infected female flowers and infected asexual flowers were compared with respect to locule tissues using periodic acid-Schiff stain. The images are of cross-sections through an anther locule. a**–e** Anther tissues of healthy male flowers at stages II to VI; **f–j** single locule of healthy male flowers at stages II to VI; k**–o** single locule of infected male flowers at stages II to VI; **p–t** single locule of infected female flowers at stages II to VI;, **u–y** single locule of infected asexual flowers at stages II to VI. Arrows indicate smut fungus. Archexporial (Ar), abnormal tapetum (At), different morphology tapetum (Dt), epidermal cells (E), endocecium (En), inner secondary parietal cells (ISP), middle layer (M), outer secondary parietal cells (OSP), primary parietal cells (PP), primary sporogeneous cells (Ps), sporogenous cells (Sp), tapetum (T). **a**, **e**, **I**, **m**, **o**, bars = 50 μm; **b–d**, **f–h**, **j–l**, **r–t**, bars = 100 μm.

**Fig. 5.**
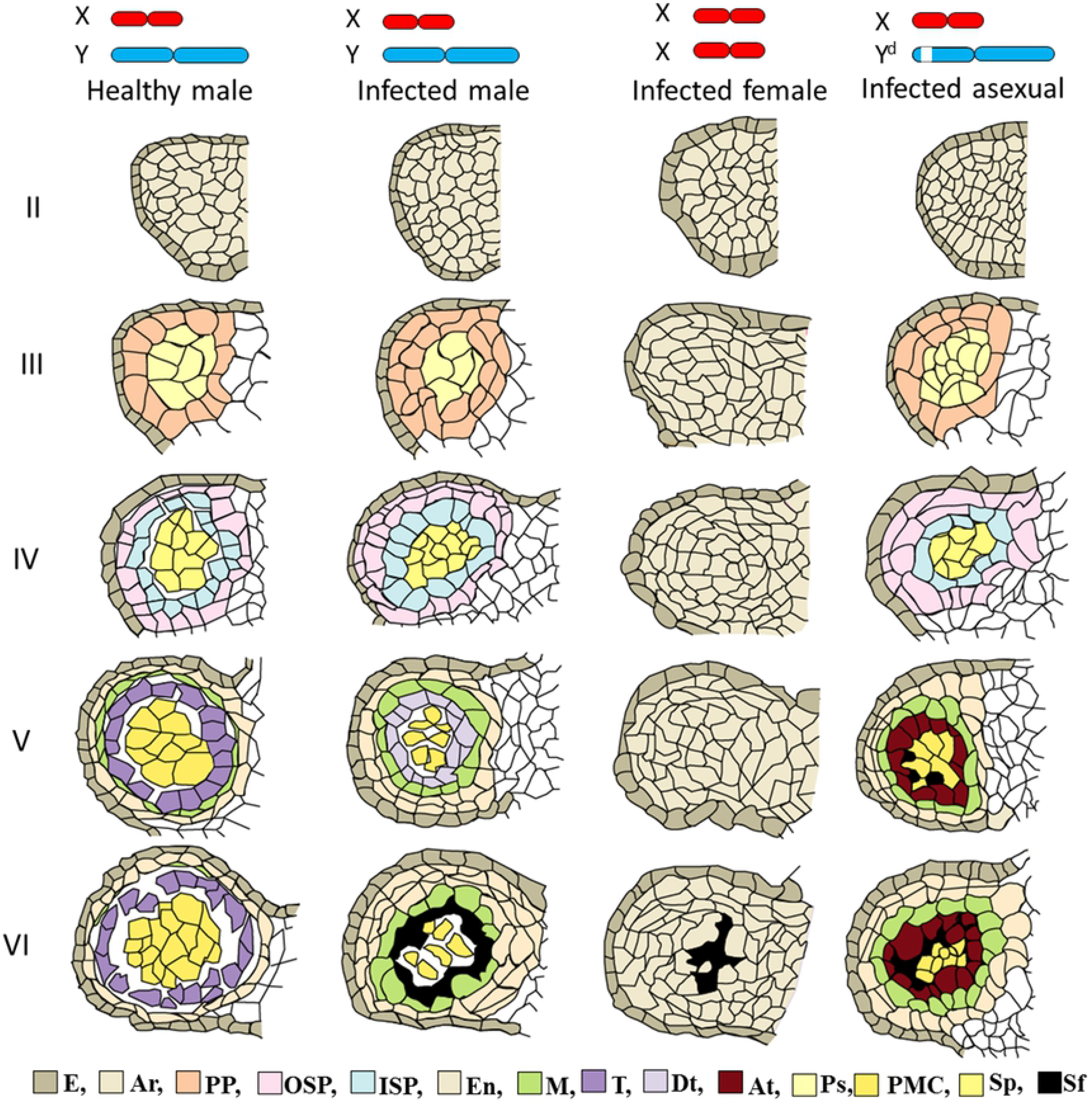
Schematic diagram of anther development in healthy male flowers, infected male flowers, infected female flowers, and infected asexual flowers. Tapetum and sporogenous tissues are presented in the healthy male flower and infected male flowers. Tapetum and sporogenous tissues are absent in infected female flowers. Sporogenous tissues are present in infected asexual flowers, but tapetum is absent. Archexporial cells (Ar), abnormal tapetum (At), different morphology tapetum (Dt), epidermal cells (E), endocecium cells (En), inner secondary parietal cells (ISP), middle layer (M), outer secondary parietal cells (OSP), pollen mother cells (PMC), primary parietal cells (PP), primary sporogeneous cells (Ps), sporogenous cells (Sp), smut fungus (Sf), tapetum (T).

## Discussion

### M. lychnidis-dioicae effects a change in anther developmental stages

Schematic diagrams of the developmental stage of the anther locule are shown in Fig. 5 **and S2 Fig**. Upon infection of the male flower of *S. latifolia* with *M. lychnidis-dioicae*, the endothecium was multi-layered, and the tapetum and middle layer showed different morphology from that of the healthy male. However, the endothecium, middle layer, tapetum, and pollen mother cell in the anther locule of the infected male were present as in the healthy male. The stamen of the female of *S. latifolia* is elongated by infection with *M. lychnidis-dioicae*. The anther of the female has only archesporial cells and epidermal cells in its locules, and does not cause differentiation of the archesporal cells during the development of sporogeneous cells at very early developmental stages. Therefore, we suggest that genes related to formation of the endothecium, middle layer, and tapetum are located on the Y chromosome. Farbos et al. [25] observed developmentally suppressed anthers of a healthy asexual mutant at early stages of anther development.

As a result, suppressed anthers of the asexual mutant formed archesporal cells, but parietal cells were absent. In this study, normal differentiation of the tapetum was not observed in the infected asexual mutant. Morphology of the anther locule in the infected asexual mutant resembled the *Arabidopsis dysfunctional tapetum1 (dyt1)* mutant, in which the *DYT1* gene (related to differentiation of the tapetum cells) was mutated [26]. Such genes may be lacking in the asexual mutant infected with *M. lychnidis-dioicae*. Although ESS1 and previously identified asexual mutants [25] were produced in different ways and the extent of the contribution of the *M. lychnidis-dioicae* infection to the production of abnormal tapetum cells is largely unknown, it is possible that the genes related to differentiation of the tapetum and middle layer were located around the SPF region of the ESS1 Y chromosome.

### Morphological changes induced by M. lychnidis-dioicae

It is thought that elongation of the stamen occurred in the infected female of *S. latifolia* because *M. lychnidis-dioicae* plays an alternative role in the Y chromosome of infected females of *S. latifolia*, which lack the Y chromosome [27]. *SLM2* is a homolog of the *PISTALA* (*PI*) B-class gene and is known to be expressed in the stamen and petal primordia [28]. It is thought that smut fungus regulates the expression of *SLM2* since this gene is not expressed in the stamen primordia of the female but expressed in the stamen primordia in the infected female [21]. In addition, *SLM2* is not expressed in the stamen primordia of asexual mutants, in which the anther development does not occur as in the female [15]. The asexual mutant has a pair of X and Y chromosomes. In the mutant, genes related to gynoecium suppression located in the GSF region are intact, whereas the SPF region is deleted. Therefore, it is thought that development of the stamen does not occur after stage 7 of flower development. This is due to the suppression of gynoecium development caused by GSF function and a lack of promotion of stamen development due to the deletion of the SPF region. Development of the gynoecium is suppressed and replaced by the filamentous structure because the GSF on the Y chromosome is intact (Fig. 1; Fig. 2). When *M. lychnidis-dioicae* infects the asexual mutant, the gynoecium displays a filamentous structure as in the healthy asexual flower mutant. Therefore, it is suggested that the suppression of gynoecium development was not released by the infection of *M. lychnidis-dioicae*. In addition, it is thought that *M. lychnidis-dioicae* has a function similar to SPF since the elongation of the stamen that is not observed in the healthy asexual mutant was observed after stage 8 of flower bud development.

### M. lychnidis-dioicae localization and timing of infection influence anther locules

*M. lychnidis-dioicae* was observed among vascular bundles, inter cellular regions of epidermal cells, root tip cells, and apical cells after infection [8]. As a result, it is thought that *M. lychnidis-dioicae* penetrated into the anther through the vascular bundles, and that the timing of *M. lychnidis-dioicae* infection was at stage 7 of flower bud development. This is because M. lychnidis-dioicae was observed in connective cells at stage III of anther development, which corresponds to stage 7 of flower bud development, and development of the stamen did not occur at stage 8 of flower bud development (Fig. 3). The growth area of *M. lychnidis-dioicae* in the anther locule is different among the infected male, the infected asexual mutant, and the infected female (Fig. 4 n, s, x). We suggest that *M. lychnidis-dioicae* was able to recognize the tapetum, such that the growth of *M. lychnidis-dioicae* started with tapetum dissolution and was observed at the position where the tapetum cells were present (Fig. 4 x). Therefore, we suggest that the different growth areas of *M. lychnidis-dioicae* observed in infected males, infected asexual mutants and infected females was due to the absence of the tapetum.

## Acknowledgements

We thank Dr. Micheal E Hood for his generous gift of *M. lychnidis-dioicae*. This work was supported by a Grant-in-Aid for Exploratory Research (to SK 24657046) from the Ministry of Education, Culture, Sports, Science and Technology, Japan. In addition, this work was supported by a Grant-in-Aid for the promotion of science from RIKEN.

## Supporting information

**S1 Fig. Scanning electron microphotographs of S. latifolia infected female flower buds development.Infected female flower buds are shown partially dissected to reveal internal structures at successive stages of development. Developmental stages of the infected asexual flowers are divided into 12 stages in this study (a-n)**. Stages 1 to 7 are shown in the tranverse view; remaining stages are shown in the longitudinal view. Early stages are observed bisexual (a-f), and later stages are observed pistil development (g-l). a Infected female flowers at stage1, b Infected female flowers at stage2, c Infected female flowers at stage3, d Infected female flowers at stage4, e Infected female flowers at stage5, f Infected female flowers at stage6, g Infected female flowers at stage7, h Infected female flowers at stage8, i Infected female anthers at stage9,j Infected female anthers at stage10, k Infected female flowers at stage11, l Infected female flowers at satge12, The inset bright field microphotograph in lshows an open flower. Anthers (an), Sepal (se), Stamen (st), Style (Sty), Gynoecium (g), Petal (p), and filaments (f), Bar = 500 μm, Double-bar = 1cm

**S2 Fig. Schematic diagram of the anther cell fate in healthy male flowers, infected male flowers, infected female flowers, and infected asexual flowers.** Archexporial cells (Ar), Abnormal tapetum (At), Different morphology tapetum (Dt), Epidermal cells (E), Endocecium cells (En), Inner secondary parietal cells (ISP), Middle layer (M), microspore (Ms),Outer secondary parietal cells (OSP), Pollen mother cells (PMC), Primaryparietal cells(PP), Primary sporogeneous cells (Ps), Sporogenous cells (Sp), Smut fungus hyphae (Sfh), tapetum (T), Tetrads (Tds)

